# An innovative 2D-mesh model to improve deltoid moment arms, muscle forces and glenohumeral joint reaction force estimations

**DOI:** 10.1101/2022.09.30.510362

**Authors:** Marta Strzelczak, Margaux Peixoto, Lauranne Sins, Mickaël Begon, Nicola Hagemeister

## Abstract

We proposed a 2D-mesh representation of the deltoid muscle to improve a realism of forces and moments predictions in a multibody shoulder model. Our 2D-mesh model was qualitatively compared to a commonly used via-point model during an arm elevation in the scapular plane and to previous in-vitro studies in terms of deltoid moment arms. The most notable improvement was observed for the moment arms of the anterior deltoid, which shows better agreement with in-vitro studies. Consequently, joint reaction force above 90° of arm elevation was more consistent with in-vivo data, validating the 2D-mesh representation of the deltoid muscle.

## Introduction

Multibody musculoskeletal models are commonly used to estimate muscle forces. In musculoskeletal models, muscle fibers are typically modelled as elastic strings from origin to insertion points to estimate their trajectories. The trajectories can be constrained by wrapping objects, i.e. by geometrical surfaces representing underlying bone structures. Muscles with broad attachment sites, such as the deltoid, are the most challenging to represent in musculoskeletal models. The deltoid muscle is usually represented by 3 to 12 independent elastic strings (Van der Helm et al. 1992; Quental et al. 2015). Via-points (fixed nodes in a local coordinate system) are commonly used to prevent deltoid fibers sliding down the wrapping object that represents the humeral head. However, presence of the via-points may lead to non-anatomical deltoid fiber trajectories (Sins et al. 2015), especially in overhead movements (e.g. high arm elevation). In addition, most of the models assume that all fibers within a muscle deform independently of their neighboring fibers, therefore losing the muscle integrity. To reproduce muscle geometry and to account for interaction between fibers other researchers have introduced a 2D-mesh muscle model with transverse constraints between muscle fibers. The 2D-mesh muscle model was previously implemented to the deltoid (Marsden 2010) and the rotator cuff muscles (Hoffmann et al. 2017) to better predict muscle length and moment arms. The 2D-mesh representation of rotator cuff muscles presents advantages over the independent lines model in terms muscles length and moment arm estimations (Hoffmann et al. 2017). Yet the above-mentioned studies implemented the 2D-mesh model in shoulder models that considered the shoulder as a ball and socket joint. Moreover, they did not investigate the effects of 2D-mesh representation on model dynamics.

The objective of this study is to reproduce the complex deltoid geometry by incorporating a 2D-mesh representation of the muscle (Fig.2 - down) into an existing multibody musculoskeletal model from Sins et al. (2015) that allows from small gleno-humeral translations during arm elevation. In complement to previous publications, besides differences in muscle geometry and moment arms, we aim to report changes in model dynamics (muscle activation patterns, muscles’ contribution to joint force and moments and glenohumeral joint reaction force (GH-JRF)). We hypothesized that the addition of transverse constraints to the deltoid model would lead to more consistent estimations of these parameters compared to the original via-point model.

**Figure 1.**
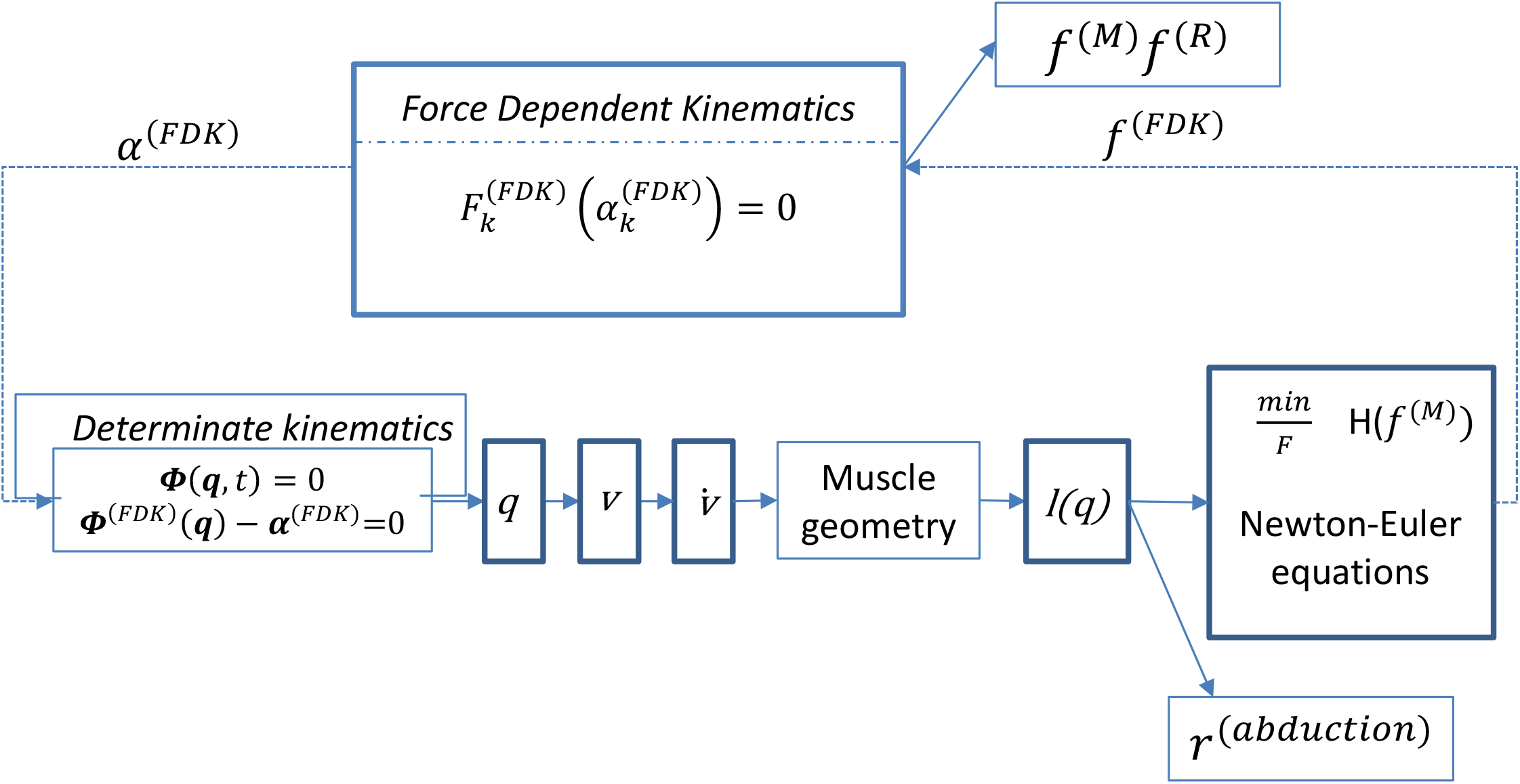
A Simplified schema of the modelling workflow in the Anybody Modelling System with integrated FDK algorithm: *α* ^*(FDK)*^ is the humeral head translation; *q* is a vector containing the position vectors and Euler parameter of all segments; *v* is a vector containing the linear and angular velocity vectors of all segments; (v) is a vector containing the linear and angular acceleration vectors of all segments; *l* is a muscle length, *H(f* ^*m*^) is the objective function, which is expressed as a function of the muscle forces *f*^*(M)*^; *r* ^*(abduction)*^ is an abduction muscle moment arm; *f* ^*(FDK)*^ is the FDK residual force for a given *α* ^*(FDK)*^; *f* ^*(M*)^ are calculated muscle forces; f ^(R)^ are calculated joint reaction forces.

**Figure 2.**
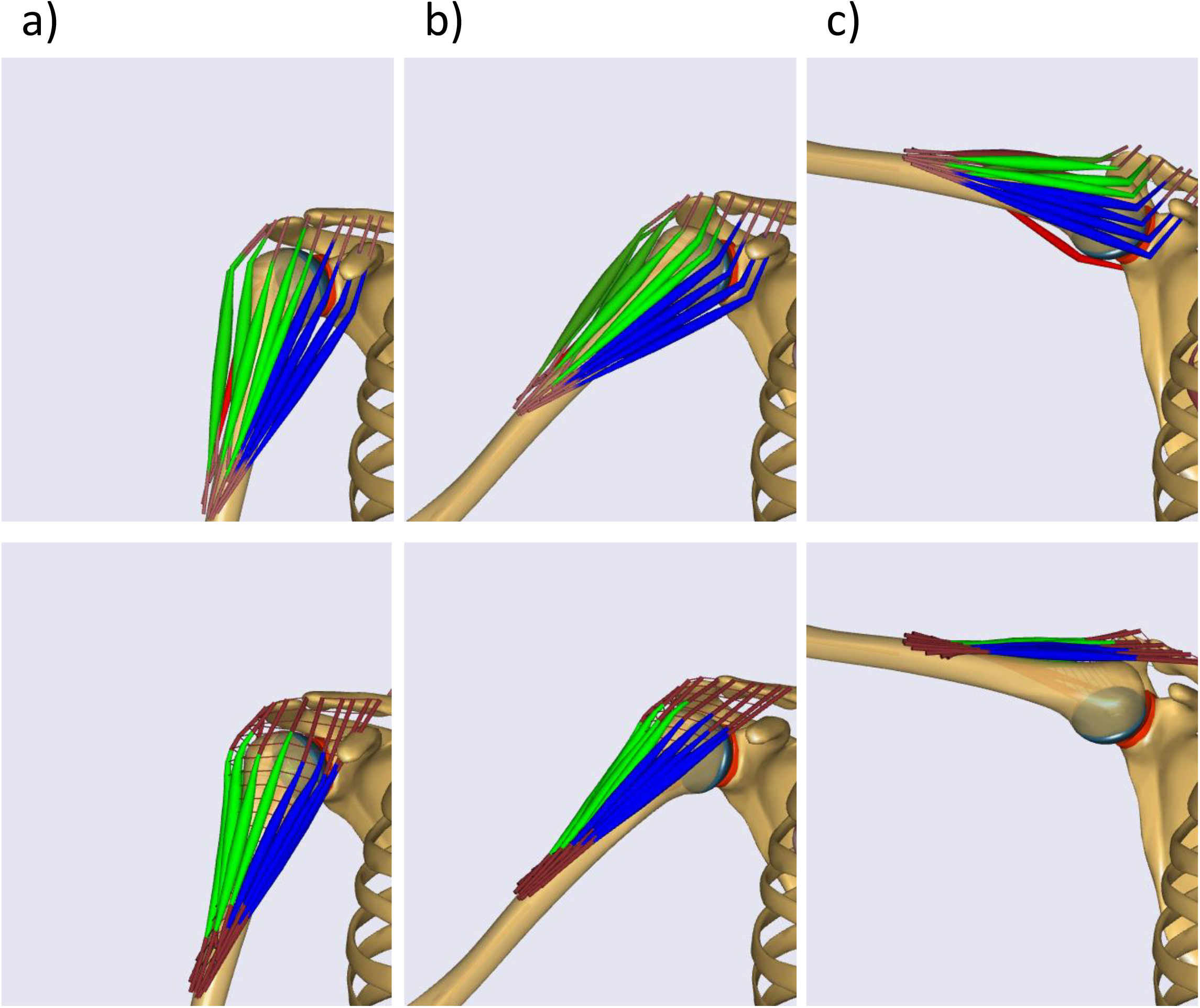
The via-point (up) and the 2D-mesh (down) deltoid models in a) 15°, b) 45° and c) 100° of arm elevation in the scapular plane. Fibres of anterior deltoid are shown in blue, lateral in green and posterior in red. Please notice that other muscles have been hidden to better show changes in the deltoid geometry.

## Methods

A multibody musculoskeletal upper-limb model was previously developed using the Anybody Modelling System (Anybody Technology, Aalborg, DK) (Sins et al. 2015). The upper-limb is modelled with 12 degrees-of-freedom (dof) namely: sternoclavicular (3-dof), glenohumeral (3-dof in rotation and 2-dof in translation), elbow (2-dof), and wrist (2-dof) joints. Elbow and wrist joints are locked in extended position. The glenohumeral joint is replaced by a non-conforming prosthesis where the anteroposterior (AP) and the inferior-superior (IS) joint translations (α^(FDK)^) are modelled using the force-dependent kinematics (FDK) (Andersen et al. 2017). Briefly, the FDK takes into account the joint elasticity and geometry and is based on quasi-static equilibrium between forces (F^(FDK)^) in the IS and AP directions α^(FDK)^ (Sins et al. 2015; Andersen et al. 2017). Therefore, this shoulder model is no longer a ball and socket representation of the glenohumeral joint. The model contains 118 Hill-type muscle-fiber elements including intact rotator cuff and deltoid muscle. The original deltoid model is composed of 12 independent lines of action (5 for the anterior, 5 for the medial, and 2 for the posterior deltoid) (Fig. 2-up). A spherical wrapping object (36-mm radius), centered in the humeral head, is used to prevent penetration of the fibers into the bone and thus to reproduce the curved deltoid fibers’ trajectory. The muscle path is computed as the shortest path from the origin to the insertion. Additionally, 12 via-points, one for each muscle fiber element, are rigidly fixed to a supplementary phantom segment. Via-points are defined to maintain muscle fibers trajectories and to prevent their uncontrolled sliding on both sides of the humeral head when searching for the shortest path.

In the proposed 2D-mesh model, the inbuilt sheet muscles from the Anybody Modelling System (Damsgaard et al. 2006) is used to represent the deltoid muscle (Fig. 2-down). The via-points from the original model are removed and each muscle-fiber element is composed of a series of 21 longitudinal linear springs connected by nodes. Each node is also connected with transverse linear spring with a node located on the neighboring muscle fiber, altogether forming a 2D-mesh. The stiffness of longitudinal and transverse springs is defined by a longitudinal : transverse ratio of 10:1 for all compartments, which is in line with the Young modulus of Morrow, Haut Donahue, Odegard, & Kaufman (2010). The ratio is confirmed using a sensitivity analysis (unpublished data). To maintain the bulky shape of the deltoid muscle longitudinal stiffness has to be at least 10 times higher than transverse stiffness. The nodes and longitudinal springs are in contact with the same glenohumeral wrapping sphere as described in the original model. The contact between each node and the wrapping sphere is governed by a static equilibrium of minimum potential elastic energy of the mesh.

An arm elevation in the scapular plane is simulated from 15° to 120° for one male subject (84 kg) using a Fourier sine expansion as a driver function in abduction (Eq. (1)), where θ is a position (rad) of the humerus relative to the thorax. The hand is constrained to stay in the scapular plane. Scapular and clavicular movements are simulated based on linear regression equations proposed by (De Groot & Brand 2001).

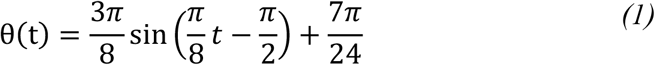

Both models are based on the inverse dynamics approach (Fig.1). The solution of muscle recruitment is formulated as an optimization problem, where a cost function has been defined as the second-order polynomial criterion, in which a square sum of muscle strength is minimized (Damsgaard et al. 2006). Anterior, lateral and posterior deltoid moment arms, activation patterns, contribution of all shoulder muscles to glenohumeral joint moments and to GHJ-RF as well as resultant GHJ-RF are estimated using 1) the original via-point model and 2) the 2D-mesh model for qualitative comparison.

## Results

### Visual analysis of deltoid muscle shape in the scapular plane elevation

In the original via-point model, the anterior fibers adopt a V-shape around 45° of arm elevation. The lateral fibers adopt a V-shape around 90° of arm elevation. In the 2D-mesh model this V-shape is no longer present (Fig. 2).

### Deltoid moment arms estimation and activation patterns

The most notable differences in terms of muscle moment arms and muscle activation are observed for the anterior deltoid (Fig.3). In the via-point model, the anterior deltoid indicates adduction function. On the contrary, in the 2D mesh model, the anterior deltoid indicates abduction function (Fig. 3a - left). Consequently, in the via-point model, the anterior deltoid remains inactive during the entire elevation movement, which explains the lack of muscle force produced by the anterior deltoid. In the 2D-mesh model, the anterior deltoid is activated around 40° of elevation (Fig. 3b - left)

**Figure 3.**
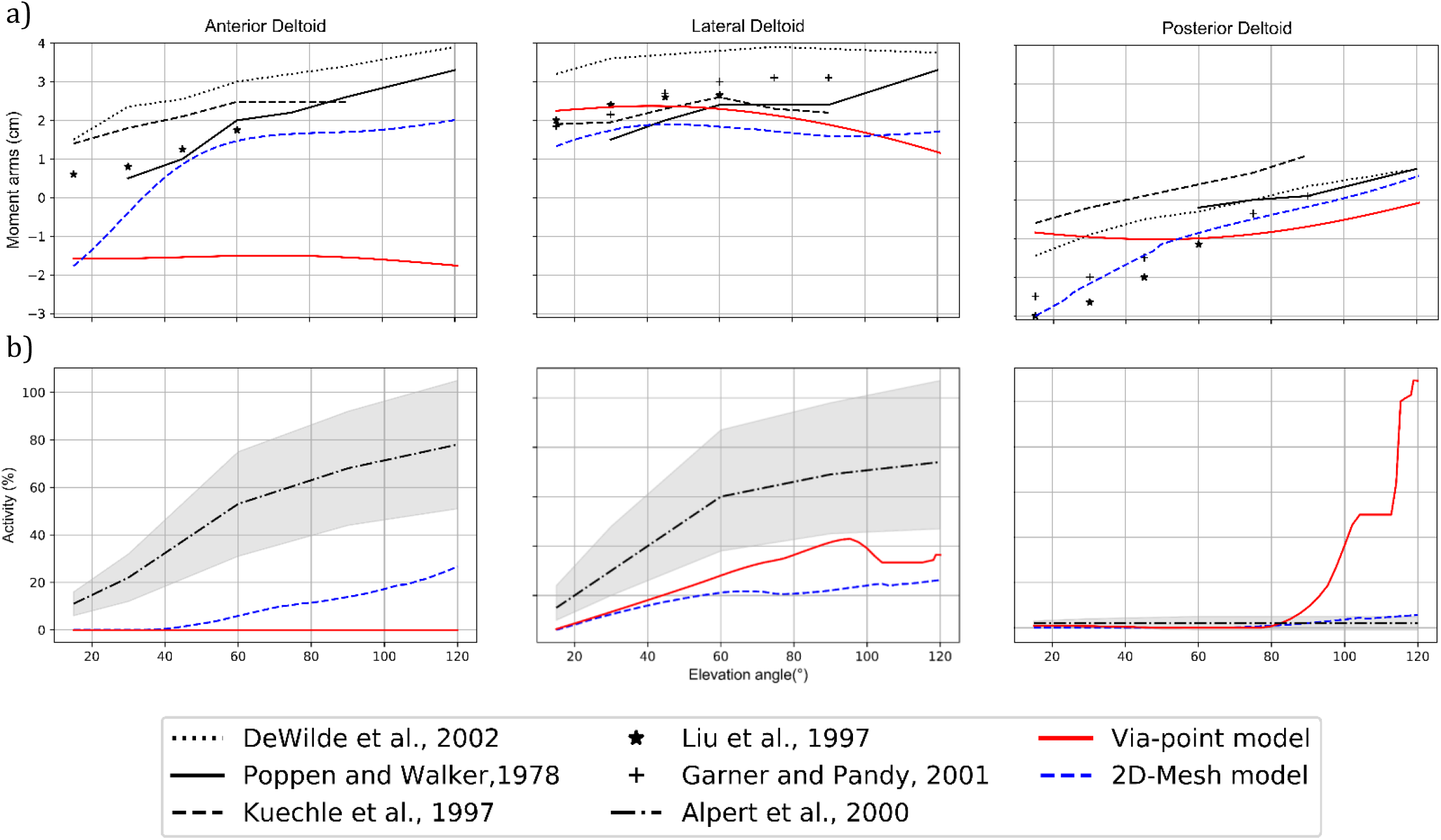
a) Predicted left-anterior, middle-lateral, right-posterior deltoid moment arms model during an arm elevation in the scapular plane using the via-point (red) and the 2D-mesh (blue) model juxtaposed with literature data. b) Predicted deltoid muscle activation using the via-point (red) and the 2D-mesh (blue) model juxtaposed with normalized EMG (black) with standard deviation (grey area) from Alpert et al., 2000.

### Contribution of all shoulder muscles to glenohumeral joint moments

In the via-point model, mainly the lateral deltoid and the infraspinatus contribute to the joint abduction moment above 90° of arm elevation (Fig. 4 - left). The lateral and the posterior chiefs of the deltoid as well as the infraspinatus generate the moment in external rotation. High external rotation moment is compensated by the subscapularis and the coracobrachialis (Fig. 4b - left). Further, the subscapularis activation results in undesired adduction moment that resists the arm elevation (Fig. 4a - left). In the 2D-mesh model, the lateral deltoid contributes to the abduction moment arm throughout the entire elevation movement with the help of the anterior deltoid at higher elevation angles (Fig. 4a - right). The infraspinatus is much less involved, meaning that the subscapularis does not counterbalance in rotation and does not produce resistance in abduction.

**Figure 4.**
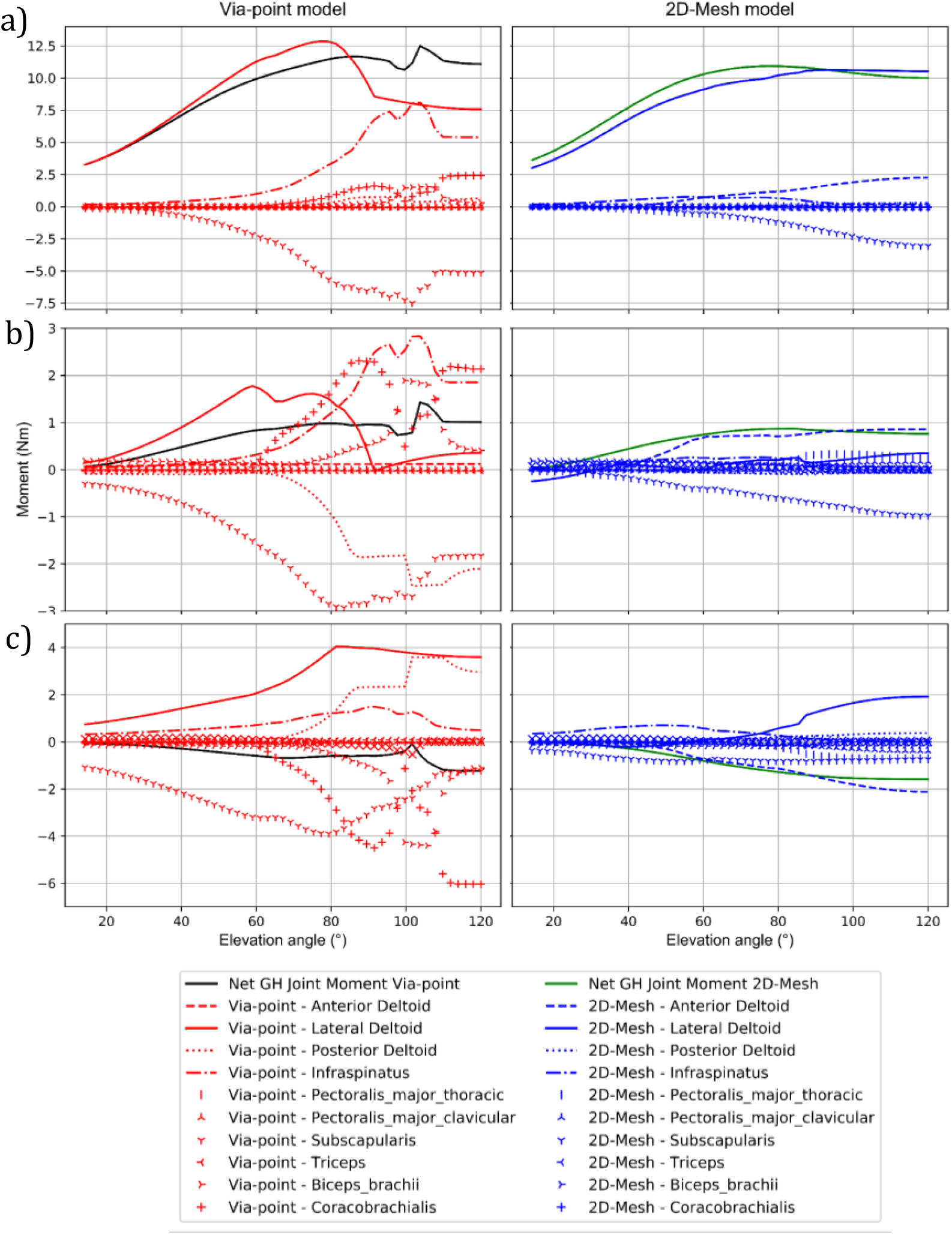
Predicted muscles contribution to joint a) abduction, b) rotation and c) flexion moments by the via-point (left) and the 2D-mesh (right) model during an arm elevation in the scapular plane. for more complex movements

### Contribution of muscle forces and resultant GH-JRF

In the via-point model more muscles with higher force magnitude contribute to GH-JRF than in the 2D-mesh model. The biggest differences between models are observed for the infraspinatus (Fig. 5 - left). Consequently, changes in the individual muscle contributions lead to differences in predicted GHJ-RF between models at high elevation angle (Fig. 5 - right). In the via-point model, the GHJ-RF is in line with a telemetric prosthesis study (Bergmann et al. 2011) until 80 ° of arm elevation. Above 80°, GHJ-RF suddenly increases reaching 1134 N around 105°. In the 2D-mesh model, the predicted GHJ-RF follows the trend reported in the (Bergmann et al. 2011) during the entire arm elevation.

**Figure 5.**
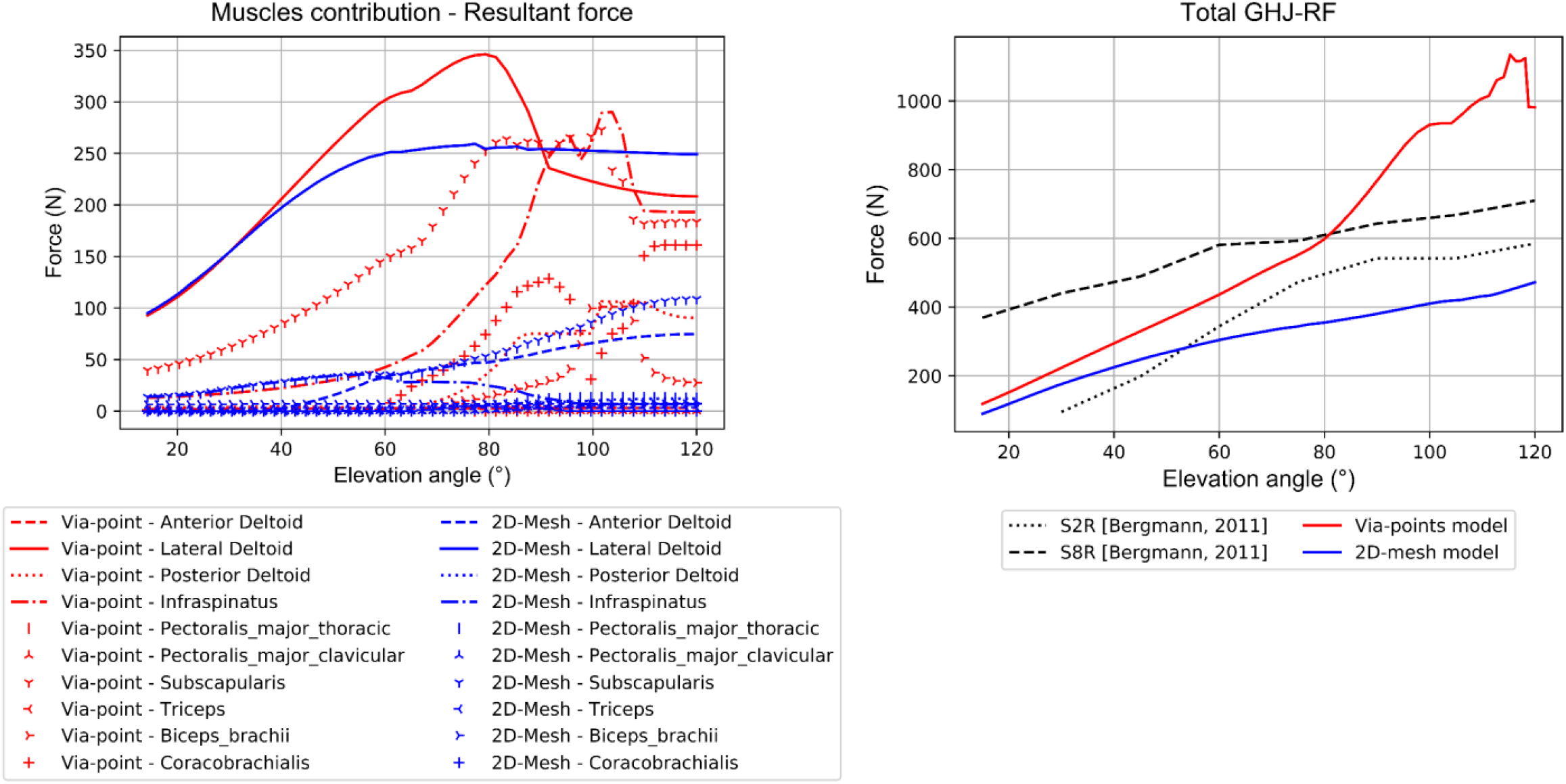
Left – Predicted resultant force for upper limb muscles acting on the glenohumeral joint by the via-point (red) and the 2D-mesh (blue) model during an arm elevation in the scapular plane. Right – Glenohumeral joint reaction force (GHJ-RF) for the via-point (red) and the 2D-mesh (blue) model in comparison with in-vivo data from telemetric prosthesis.

## Discussion

In the present study, we modelled the deltoid muscle as a 2D-mesh in the multibody musculoskeletal shoulder model to improve its geometry and assess the effect on model dynamics. The 2D-mesh representation was successfully implemented to shoulder muscles with broad attachment sites (Marsden 2010; Hoffmann et al. 2017). Nevertheless, to our knowledge, we are the first to examine impact of 2D-mesh representation on deltoid muscle activation, individual muscle contribution to GHJ-RF and moments and resultant GHJ-RF. Additionally, previous models represented the glenohumeral articulation as a simplified ball-and-socket joint with 3-dof. In the present study the glenohumeral articulation was modelled with 5-dof, which more accurately reflects the anatomical joint. Moreover, unlike in the previous studies, the multibody shoulder model (Sins et al., 2015) is accessible to the modelling community via the Anybody Modelling System (Damsgaard et al. 2006).

As expected, the 2D-mesh model showed improvements in reproducing the muscle geometry. In fact, by integrating transverse constrains into the deltoid model, the rounded contour of deltoid muscle fibers overlying humeral head was maintained, whereas in the via-point model anterior fibers adopt a V-shape around 45° of arm elevation and lateral fibers adopt a V-shape around 90°. The V-shape has occurred due to the presence of rigidly fixed via-points, which constrain the fibers’ path. Simply removing the via-points would result in sliding of the fibers anteriorly or posteriorly because of the shortest path algorithm that is used.

Tendon excursion studies using cadaveric models indicated abduction moment arms for the anterior deltoid, which increase progressively with increasing elevation (Kuechle et al. 1997; Liu et al. 1997). The via-point model showed adduction moment arms during whole elevation movement, meaning that in the original model, the anterior deltoid acted solely as an adductor. The 2D-mesh model correctly predicted an abduction moment arms with a similar trend as experimental studies above 35°(Kuechle et al. 1997; Liu et al. 1997). However, below 35°, the 2D-mesh model predicted adduction moment arms and at these low elevation angles, we observed a penetration of anterior fibers into the coracoid process. Bone penetration might indicate that the model would need an extra constraint such as an additional wrapping surface.

The 2D-mesh solved the problem of non-anatomical fibers trajectories and misleading anterior deltoid function related to predicted moment arms. However, discrepancies in the existing literature (Kuechle et al. 1997; Liu et al. 1997; De Wilde et al. 2002) suggest that more subject specific validation should be used to assess further development of 2D-mesh model. Geometrical representation of the deltoid muscle affected muscle activation and muscle force predictions. Important differences, especially in anterior deltoid activation, have been found between numerical and experimental (based on electromyography (EMG)) studies. Only onset and offset of muscles activation as well as trends could be compared between models prediction and EMG data (van der Helm 1994). Indeed, some musculoskeletal models predict no substantial activation of the anterior deltoid during the arm abduction in the scapular plane (Dickerson et al. 2008; Sins et al. 2015) while experimental studies reported muscle activation up to 78 % of maximum muscle contraction (Alpert et al. 2000) in high elevation angles. The 2D-mesh largely solved the problem of lack of activation in the anterior deltoid.

Changes in deltoid muscle geometry affected not only deltoid activation, but also the overall muscles’ recruitment pattern. In the via-point model, several shoulder muscles generated high joint moments, which were above or opposite to the net joint moment. In the via-point model, other muscles, especially the infraspinatus, compensated for the lack of abduction moment capacity produced by the anterior deltoid. This results in a series of muscle activation to ensure the balance of moments in all three planes. Indeed, the contraction of infraspinatus together with the lateral and the posterior deltoid generated an external rotation moment. This external rotation moment needed to be balanced by other muscles to produce a small internal rotation moment to elevate shoulder in the scapular plane. EMG studies (Townsend et al. 1991; Reddy et al. 2000; Alpert et al. 2000) reported only co-activation of the infraspinatus and the subscapularis muscles, which is in line with activation patterns of the 2D-mesh model.

Consequently, the additional activation of these muscles generated forces that affected GHJ-RF. The biggest difference in the muscles’ contribution between two models appeared above 90° of arm elevation, where the V-shape caused by the via-points is present for all deltoid fibers in the original via-point model. Therefore, the contribution of individual muscles explained differences in predicted total GHJ-RF between models. Predicted GHJ-RF by both models were compared to in vivo data obtained using telemetric prostheses from Bergmann et al. (2011). The via-point model followed the trend from Bergmann et al. (2011) study only until 90° of arm elevation. Above 90°, non-physiologically high loads in glenohumeral joint were observed. Prediction of GHJ-RF improved significantly with the 2D-mesh model. The 2D-mesh model followed the trend from Bergmann et al. (2011) study during entire elevation movement. Even so, differences in the force magnitude was observed between 2D-mesh model and the experimental study (Bergmann et al. 2011). The underestimation may come from the contribution of the rotator cuffs muscles – intact in the model and often deficient in arthroplasty patients (Sanchez-Sotelo et al. 2001), differences in joint kinematics which was not recorded in Bergmann et al. (2011) and, above all, the cost function in the static optimization. Least quadratic activation function is known to underestimate muscles forces and co-contraction especially in the shoulder (Forster et al. 2004) resulting in lower joint contact forces.

There is still space for improvement in the development of a 2D-mesh model. We evaluated 2D-mesh model only for scapular plane elevation, since it is the most commonly reported movement in the literature. To assess further the finding of this study, model needs to be evaluated during the full range of motion. However, there is a lack of experimental data to validate musculoskeletal models of the shoulder during more complex movements and larger ranges of motion.

## Conclusion

The 2D-mesh deltoid model presents more consistent predictions in terms of shoulder dynamics than the model with independent lines of action and rigidly fixed via-points. The 2D-mesh shows the ability to maintaining correct muscle shape even at high elevation. Besides that, the 2D-mesh deltoid model demonstrates more realistic estimations of deltoid muscle activation, more realistic muscle recruitment patterns and better prediction of GHJ-RF when compared with the literature. This study shows that the geometrical representation of the deltoid considerably affects model predictions. Incorrect representation of deltoid muscle geometry can lead to misleading predictions of muscle recruitment as well as joint forces and consequently, limits model applications or leads to wrong interpretations.

## Acknowledgments

None

## References

Alpert SW, Pink MM, Jobe FW, McMahon PJ, Mathiyakom W. 2000. Electromyographic analysis of deltoid and rotator cuff function under varying loads and speeds. J shoulder Elb Surg. 9(1):47–58. 10.1103/PhysRevE.77.031608

Andersen MS, De Zee M, Damsgaard M, Nolte D, Rasmussen J. 2017. Introduction to force-dependent kinematics: Theory and application to mandible modeling. J Biomech Eng. 139(9):091001. 10.1115/1.4037100

Bergmann G, Graichen F, Bender A, Rohlmann A, Halder A, Beier A, Westerhoff P. 2011. In vivo gleno-humeral joint loads during forward flexion and abduction. J Biomech. 44(8):1543–1552. 10.1016/j.jbiomech.2011.02.142

Damsgaard M, Rasmussen J, Christensen ST, Surma E, de Zee M. 2006. Analysis of musculoskeletal systems in the AnyBody Modeling System. Simul Model Pract Theory. 14(8):1100–1111. 10.1016/j.simpat.2006.09.001

Dickerson CR, Hughes RE, Chaffin DB. 2008. Experimental evaluation of a computational shoulder musculoskeletal model. Clin Biomech. 23(7):886–894. 10.1016/j.clinbiomech.2008.04.004

Forster E, Simon U, Augat P, Claes L. 2004. Extension of a state-of-the-art optimization criterion to predict co-contraction. J Biomech. 37(4):577–581. 10.1016/j.jbiomech.2003.09.003

De Groot JH, Brand R. 2001. A three-dimensional regression model of the shoulder rhythm. Clin Biomech. 16(9):735–743. 10.1016/S0268-0033(01)00065-1

van der Helm FCT. 1994. A finite element musculoskeletal model of the shoulder mechanism. J Biomech. 27(5):551–569. 10.1016/0021-9290(94)90065-5

Van der Helm FCT, Veeger HEJ, Pronk GM, Van der Woude LHV, Rozendal RH. 1992. Geometry parameters for musculoskeletal modelling of the shoulder system. J Biomech. 25(2):129–144. 10.1016/0021-9290(92)90270-B

Hoffmann M, Haering D, Begon M. 2017. Comparison between line and surface mesh models to represent the rotator cuff muscle geometry in musculoskeletal models. Comput Methods Biomech Biomed Engin. 20(11):1175–1181. 10.1080/10255842.2017.1340463

Kuechle DK, Newman SR, Itoi E, Morrey BF, An KN. 1997. Shoulder muscle moment arms during horizontal flexion and elevation. J Shoulder Elbow Surg. 6(5):429–439. 10.1016/S1058-2746(97)70049-1

Liu J, Hughes R, Smutz W, Niebur G, Nan-An K. 1997. Roles of deltoid and rotator cuff muscles in shoulder elevation. Clin Biomech. 12(1):32–38. 10.1016/S0268-0033(96)00047-2

Marsden S. 2010. Muscle Wrapping Techniques Applied to the Shoulder [dissertation]. Newcastle Upon Tyne (UK): University of Newcastle upon Tyne.

Morrow DA, Haut Donahue TL, Odegard GM, Kaufman KR. 2010. Transversely isotropic tensile material properties of skeletal muscle tissue. J Mech Behav Biomed Mater. 3(1):124–129. 10.1016/j.jmbbm.2009.03.004

Quental C, Folgado J, Ambrósio J, Monteiro J. 2015. Critical analysis of musculoskeletal modelling complexity in multibody biomechanical models of the upper limb. Comput Methods Biomech Biomed Engin. 18(7):749–59. 10.1080/10255842.2013.845879

Reddy AS, Mohr KJ, Pink MM, Jobe FW. 2000. Electromyographic analysis of the deltoid and rotator cuff muscles in persons with subacromial impingement. J Shoulder Elb Surg. 9(6):519–523. 10.1067/mse.2000.109410

Sanchez-Sotelo J, Cofield RH, Rowland CM. 2001. Shoulder hemiarthroplasty for glenohumeral arthritis associated with severe rotator cuff deficiency. J Bone Jt Surg - Ser A. 83(12):1814–1822. 10.2106/00004623-200112000-00008

Sins L, Tétreault P, Hagemeister N, Nuño N. 2015. Adaptation of the AnyBody™ Musculoskeletal Shoulder Model to the Nonconforming Total Shoulder Arthroplasty Context. J Biomech Eng. 137(10):101006. 10.1115/1.4031330

Townsend H, Jobe FW, Pink M, Perry J. 1991. Electromyographic analysis of the glenohumeral muscles during a baseball rehabilitation program. Am J Sports Med. 19(3):264–272. 10.1177/036354659101900309

De Wilde L, Audenaert E, Barbaix E, Audenaert A, Soudan K. 2002. Consequences of deltoid muscle elongation on deltoid muscle performance: A computerised study. Clin Biomech. 17(7):499–505. 10.1016/S0268-0033(02)00065-7

